# Information transfer and recovery for the sense of touch

**DOI:** 10.1101/2020.12.08.415729

**Authors:** Chao Huang, Bernhard Englitz, Andrey Reznik, Fleur Zeldenrust, Tansu Celikel

## Abstract

Transformation of postsynaptic potentials (PSPs) into action potentials (APs) is the rate-limiting step of communication in neural networks. The efficiency of this intracellular information transfer also powerfully shapes stimulus representations in sensory cortices. Using whole-cell recordings and information-theoretic measures, we show herein that somatic PSPs accurately represent stimulus location on a trial-by-trial basis in single neurons even 4 synapses away from the sensory periphery in the whisker system. This information is largely lost during AP generation but can be rapidly (<20 ms) recovered using complementary information in local populations in a cell-type-specific manner. These results show that as sensory information is transferred from one neural locus to another, the circuits reconstruct the stimulus with high fidelity so that sensory representations of single neurons faithfully represent the stimulus in the periphery, but only in their PSPs, resulting in lossless information processing for the sense of touch in the primary somatosensory cortex.

## Introduction

Neural information processing requires signal transformation every time the information is transferred from one neuron to another. This transformation is performed in postsynaptic neurons by integrating spatiotemporally distributed synaptic inputs and generating action potentials, which then propagate information across synaptically coupled neurons. For each processing step, how much information is retained, how much of it is transferred to a postsynaptic neuron, how much is lost, and whether local networks can fully recover the lost information during this intracellular sub-to-suprathreshold information transfer are questions that have yet to be answered. In an accompanying paper (Zeldenrust et al., 2020) we show on a single neuron level that how much information is lost during action potential generation depends on the cell-class. On a network level, the effectiveness of this input-to-spike operation depends on the connectivity, the code used between the sender (i.e. presynaptic neurons), and the receiver (i.e. postsynaptic neurons) as well as the noise characteristics of the channel. Since many of these are currently impossible to assess experimentally, the rules of information transfer in biological circuits, with the exception of cell-type-specific intracellular information transfer in single neurons as outlined in the accompanying article (Zeldenrust et al., 2020), are still largely unknown.

Sensory systems in particular offer unique opportunities to study information processing in neural circuits. If the primary function of a sensory circuit is to faithfully and reliably represent the environment (Azarfar et al., 2018; DeCharms and Zador, 2000; Diamond et al., 1999; Knudsen et al., 1987), a substantial part of the sensory information in the periphery should be represented throughout the sensory circuits in the form of neural signals. Sensory systems are commonly organized in the form of topographical maps, where sensory receptors in the periphery are represented by topographically organized groups of neurons along the sensory axis (Harding-Forrester and Feldman, 2018; Kole et al., 2018; Petersen, 2019). However, the functional role of these topographical maps for sensory processing is still not clear (Chklovskii and Koulakov, 2004; Diamond et al., 1999; Kaas, 1997; Weinberg, 1997). Understanding the mechanisms of information processing, transfer and recovery is particularly important in sensory circuits, as the efficacy of signal transformation should determine the extent, speed, and accuracy of sensory representations.

Stimulating single neurons in the sensory and motor cortices can result in observable behavioral responses such as whisker movement (Brecht et al., 2004; Doron et al., 2014; Houweling and Brecht, 2008; Voigts et al., 2008). However, single neurons carry surprisingly little information in the rate and timing of their action potentials (Alenda et al., 2010; Panzeri et al., 2001; Petersen et al., 2001; Quian Quiroga and Panzeri, 2009). Given that pooling information across simultaneously recorded neighboring neurons minimally contributes to the information carried in local populations because neighboring neurons carry largely redundant information (Petersen et al., 2002, 2001), the target postsynaptic neurons are likely to reconstruct the stimulus by spatiotemporal integration across behaviorally relevant spatial and temporal scales (Azarfar et al., 2018; Celikel and Sakmann, 2007).

Here, we performed intracellular recordings and used computational modeling to address the principles of information processing in the somatosensory cortex. Surprisingly (to us), we found that the sensory stimulus can be fully reconstructed with the information available in the *subthreshold* responses of single excitatory neurons (i.e. the recorded EPSPs in L2/3 neurons). Up to 90% of this information is lost during intracellular information transfer, i.e. when an *action potential* is generated from these subthreshold responses, in agreement with previous observations on the information content of action potentials in barrel cortical neurons (Alenda et al., 2010; Panzeri et al., 2001; Petersen et al., 2002). *In vivo* information loss is likely to exceed this value, due to background ongoing activity (Destexhe et al., 2003). Next, we assessed information recovery on the *population level* using an analysis based on bootstrapped groups of neurons recorded *in vitro*. We found that information lost during action potential generation can be fully recovered by as little as 100 neurons with a time resolution of 2-3 ms. Finally, we turned to a realistic and well-constrained simulation of a barrel column (Huang et al., 2020) to study the relation between encoding strategies in L4 and decoding strategies in L2/3 to determine the mechanisms of information recovery.

Comparing candidate encoding strategies in the L4 population, we found that a population rate code (using peri-stimulus time histograms obtained in the *in vivo* recordings) is unsuitable for information transfer in a cortical network because the trial-to-trial reliability is too low to fit the high information recovery that we found as in our experiments. Codes with higher trial-to-trial reliability in timing and rate perform substantially better, with optimal performance reached if neurons fire reliably across trials. In this case, the L4 activity can be fully decoded by small groups (~25 cells) of both excitatory and inhibitory neurons in L2/3 within ~20 ms after stimulus onset, and within a few ms after the first spike response. In summary, we show that intracellular information transfer is highly lossy, and thus potentially selective. However, by combining the limited but complementary information in the spike trains of L2/3 inhibitory and excitatory neurons, single neurons could fully reconstruct stimulus resulting in lossless representation of sensory information in their subthreshold responses.

## Results

### L2/3 single cell responses to in vivo whisker stimulation

We performed whole-cell current-clamp recordings of Layer (L) 2/3 pyramidal neurons in the juvenile rat primary somatosensory cortex, in the barrel cortical subregion under ketamine anesthesia. During these *in vivo* recordings, sensory stimulation was provided by direct stimulation of the principal and 1st order surround whiskers with a piezo stimulator in 2 directions (up-down: Fig. 1A). The cumulative synaptic input in response to these stimuli was quantified in properties of the somatic post-synaptic potential (PSP), i.e. the onset time, slope, and peak amplitude (Fig. 1B-D). Principal whisker stimulation-evoked PSPs exhibited the shortest latency, as well as the highest slope and amplitude, in comparison to PSPs evoked by the stimulation of surrounding whiskers, in agreement with previous observations (Brecht et al., 2003). Although the PSPs were highly reliable (PW: 99.8%, SW: 91.8% of trials evoked PSPs), action potentials (spikes) were sparse and unreliable, even after principal whisker deflections (PW: 6.2% (SD 8.6%); SW: 1.7% (SD 2.9%) of trials included evoked APs).

**Figure 1.**
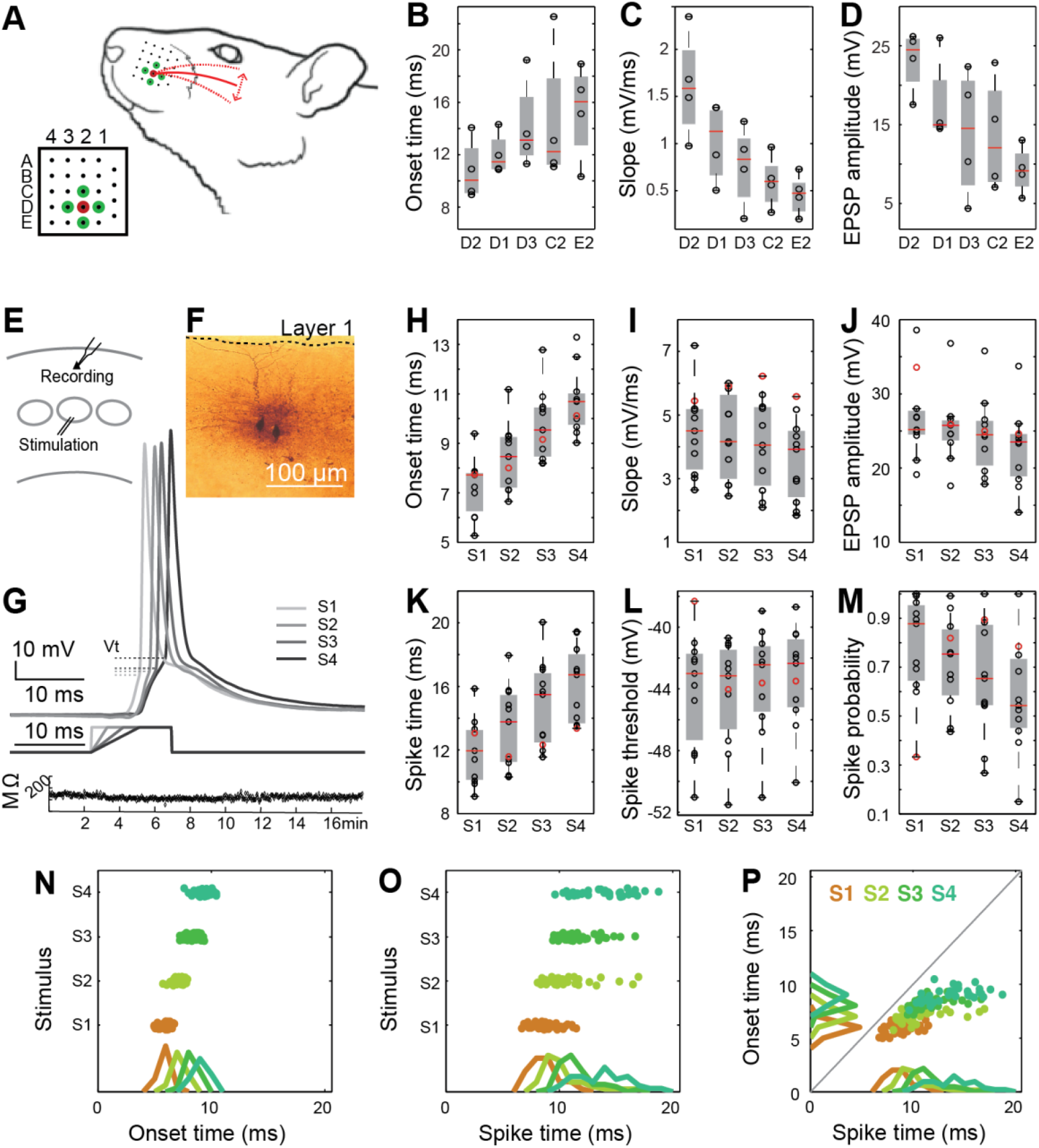
*In vivo* and *in vitro* stimulus representation in single L2/3 somatosensory cortical neurons. **(A)** We deflected whisker D2 and its first order neighbors (D1,D3,C2,E2) individually to determine the spatial encoding properties of cortical L2/3 pyramidal neurons in the D2 barrel under anesthesia using whole cell current clamp recordings. **(B-D)** EPSP response to *in vivo* stimulation. Analysis of the EPSP parameters showed that principal whisker stimulation was correlated with earlier onset times (**B**), larger slopes (**C**) and larger amplitudes (**D**) compared to the surround whiskers. Onset time was described as the latency between stimulus onset and the time it takes for the membrane to reach 10% of the peak somatic EPSP amplitude. The EPSP slope was calculated to be between 10-90% of the somatic EPSP. The amplitude was measured at the peak. All measurements were performed on monosynaptic EPSPs. **(E-M)** Response to *in vitro* stimulation mimicking *in vivo* stimulation. Due to the sparse nature of action potentials *in vivo*, we developed a stimulation protocol to mimic the subthreshold stimulus encoding properties of L2/3 neurons *in vitro.* **(E)** Whole cell intracellular current clamp recordings were performed in L2/3 while L4 neurons were stimulated using a bipolar electrode. **(F)** Soma location of randomly selected neurons. **(G)** The stimuli were direct current injections with equal maximal amplitudes as the in vivo EPSCs, but the rising slope of the current was systematically reduced across the four stimulus conditions (see (Huang et al., 2016)). **(H-M)** L2/3 pyramidal neurons’ responses to L4 stimulation. Each circle shows the average (over trials) response of one neuron (N=11). **(H-J)** EPSP response to *in vitro* stimulation. **H**: Onset time, **I**: Slope, **J**: EPSP amplitude; **(K-M)** Spike response to *in vitro* stimulation. **K**: Spike time, i.e. latency to spike after stimulus onset; **L**: Spike threshold, described as the membrane potential at which the second derivative reaches a global (positive) maximum; **M**: Action potential (i.e. spike) probability, across trials. **(N-P)** Spike versus EPSP response to in vitro stimulus. While both EPSP and spike parameters displayed an average dependence on the stimulus, EPSP parameters are more accurately determined by the stimulus than spike parameters on single trials.

### Properties of sub- & suprathreshold responses of L2/3 neurons to L4 stimulation

Mutual information calculations require long sampling durations, which limits the possibilities for unbiased calculation of information processing with high-dimensional naturalistic stimuli in vivo. Therefore we performed acute slice experiments with simplified stimuli (Fig.1E-P). Bipolar electrodes in L4 were used to deliver square pulses with varying slopes as described before (Fig.1E-F; Huang et al., (2016); see Materials and Methods). Visualized L2/3 neurons were recorded in whole-cell current clamp configuration. PSP responses of L2/3 neurons systematically varied with the four stimulus patterns (Fig. 1G-J). Spike responses showed a similar dependence on the stimulus slope when averaged over multiple trials, with delayed spike times, increased threshold and decreased spike probability for shallower slopes of stimulation (Fig. 1K-M).

The average properties shown above indicate that individual cells qualitatively correspond to whisker deflections mimicking spatial stimuli. However, during sensory processing, animals have to deduce object location from single trials, not from averages over many trials, which is only possible when trial-to-trial variability is low. Spikes exhibited a far greater temporal trial-to-trial variability than PSPs (Fig. 1N-P). PSP onset times showed an average progression with stimulus slope, with a small trial-to-trial variability (Fig. 1N, SD = 0.42-0.63 ms), whereas spike time variability was threefold higher (Fig.1O, SD = 1.2-2.3 ms). Therefore, spike times could only to a very limited degree be predicted on the basis of PSP onset times, with spikes often occurring with significant and variable delays (Fig. 1P, 2.8-4.7 (SD 1.1-2.1) ms). Spike generation was also failure prone, with average spike failures rates up to 31.7% of the trials [Range: 0-85%]. While spike failures bear information in rate-based codes, in timing-based codes information is missing when the neuron fails to fire an action potential. To study the influence of trial-to-trial variability in timing and rate on stimulus information, we calculated Shannon’s mutual information between the stimulus and several PSP and spike properties. This mutual information provides largely agnostic estimates of the transmitted information between stimulus and response.

### Information transmission between somatic PSPs and Spikes

How much information does a somatosensory neuron carry about the sensory stimulus (S) and how much of this information does it transfer to its postsynaptic targets? Surprisingly, the information between a single somatic PSP and the stimulus contains the bulk (~95%, I(S;PSP) vs. H(S), Fig. 2A) of the entropy of the sensory stimuli. The information between the onset time (81%, Fig. 2B), slope (8.4%, Fig. 2B) and amplitude (6.4%, Fig. 2B) of the PSP and the stimulus contribute largely independently to the total information content of a PSP (Fig. 2C). However most of this information is lost upon spike generation (down to 24%, I(S;(St,Vt), Fig. 2A), where spike timing (St, 16%, Fig. 2D) and voltage threshold (Vt, 6.2%, Fig. 2D) carry most of the stimulus information contained in the spikes.

**Figure 2.**
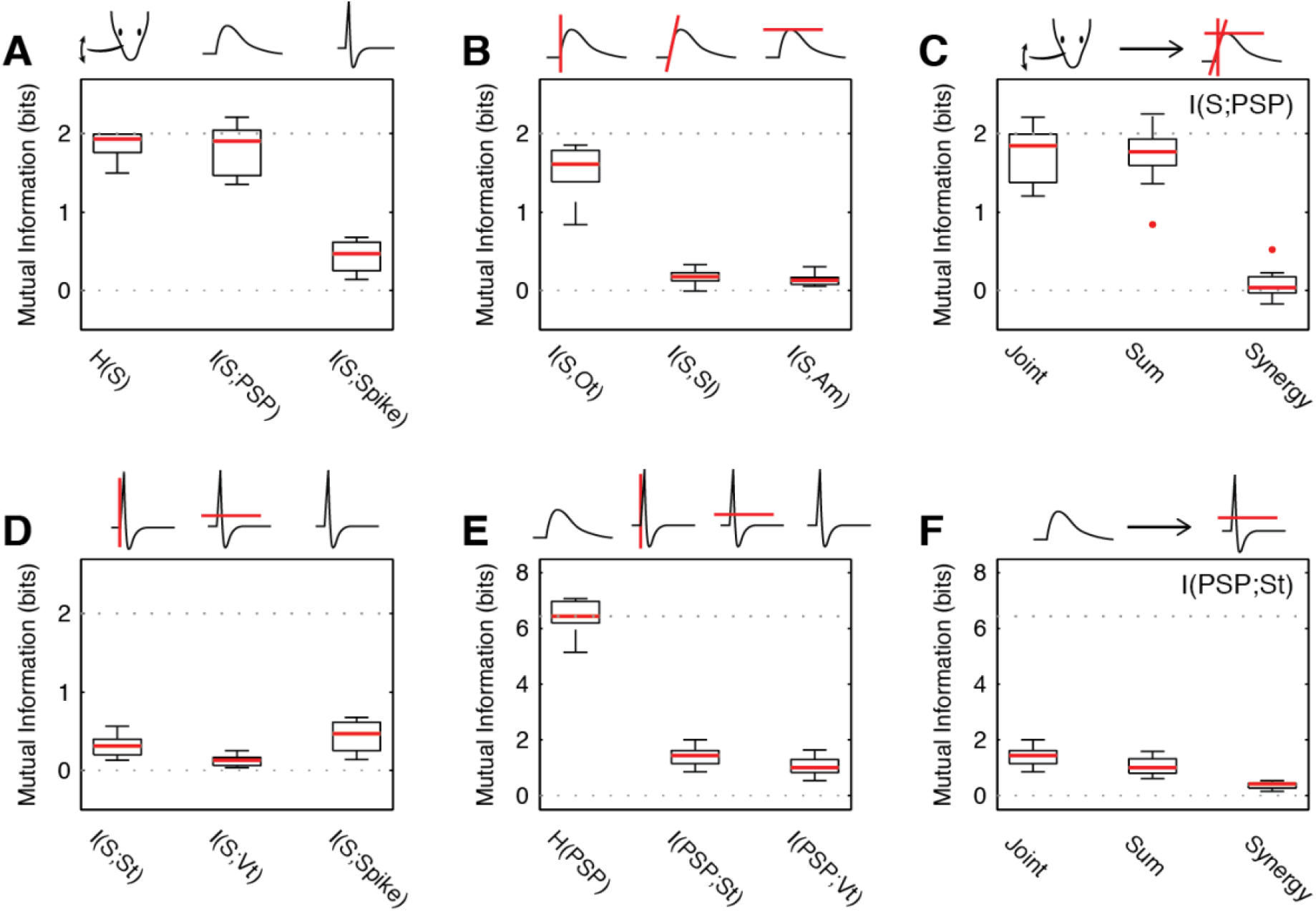
Postsynaptic potentials encode substantially more stimulus information than spikes *in vitro*. **(A)** The information between PSPs and the stimulus is significantly higher than the information between spikes and the stimulus. While the PSP contains a large fraction of the stimulus entropy (95%, I(S;PSP), 1.81±0.31 bit vs. H(S), 1.86±0.17 bit, p = 0.16), most of this information is not transferred to the spike (I(S;Spike), 0.47 ±0.19 bit, 24%). **(B)** The majority of the information in the PSP is carried by the onset timing (Ot, 1.6±0.31 bit, 85%), while slope (Sl, 0.17 ±0.09 bit, 10%) amplitude (Am, 0.13 ±0.07 bit, 5%) carry only small amounts of information. **(C)** Ot, Sl, and Am add their information independently, as the synergy between them is close to 0 (Synergy: 0.03 ±0.18 bit, p = 0.15, t-test). **(D)** The information in the spike is contributed by spike time (St, 0.31 ±0.13 bit, 16%) and threshold (Vt, 0.12 ±0.08 bit, 6.2%), and jointly only reach 21% of the total information (repeated from A). **(E)** Substantial information transfer occurs between the PSP and the spike, although this constitutes only 22% (St) and 15% (Vt) of the entropy in the PSP. **(F)** The information in the properties of the PSP adds largely independently to the joint information, with a small but highly significant synergistic contribution of different PSP properties (0.41 ±0.13 bit, 6.4%, p<10^−5^). In all figures data is plotted as inter-quartile intervals and red lines denote the median of each distribution. Outliers are plotted as red dots. The dotted line denotes the maximal stimulus entropy.

We can directly quantify how much of the information in the PSP is transferred to the spike (see Materials and Methods). Unsurprisingly, the total entropy in the PSP (i.e across onset, slope and amplitude together) exceeds the stimulus information multifold (6.4 bits, 3.2-fold, Fig. 2E). The transferred information from PSP to spike amounts to 22% and 15% of the PSP entropy for St and Vt respectively (Fig. 2E, comparison of medians). However, most of this information is redundant, since the actual amount of stimulus information contained in a spike is much lower (0.41 bit, Fig. 2A). The individual PSP properties on the other hand contribute slightly synergistically to the timing information in the spike (Fig. 2F, 6.4%, p<10^−5^). Consequently, while a substantial amount of the information about the stimulus in the PSP is transferred to the spike, this information is insufficient to encode the present stimulus space at the single neuron level on a single neuron and single trial basis.

### Information recovery in local neural populations *in silico* and *in vitro*

If the PSP-to-spike transformation causes a dramatic drop in information about the stimulus carried in the neural activity, how can the somatic PSPs of L2/3 neurons carry near complete information (Fig.2A)? Since these neurons are four synapses away from the sensory periphery, a recovery of information has to occur at the network level. Information recovery was analyzed both in an anatomically and physiologically well-constrained network model of a rat barrel column (Fig. 3) and on bootstrapped populations of the *in vitro* data (Fig. 4, see Materials and Methods and Huang et al., (2020)).

**Figure 3.**
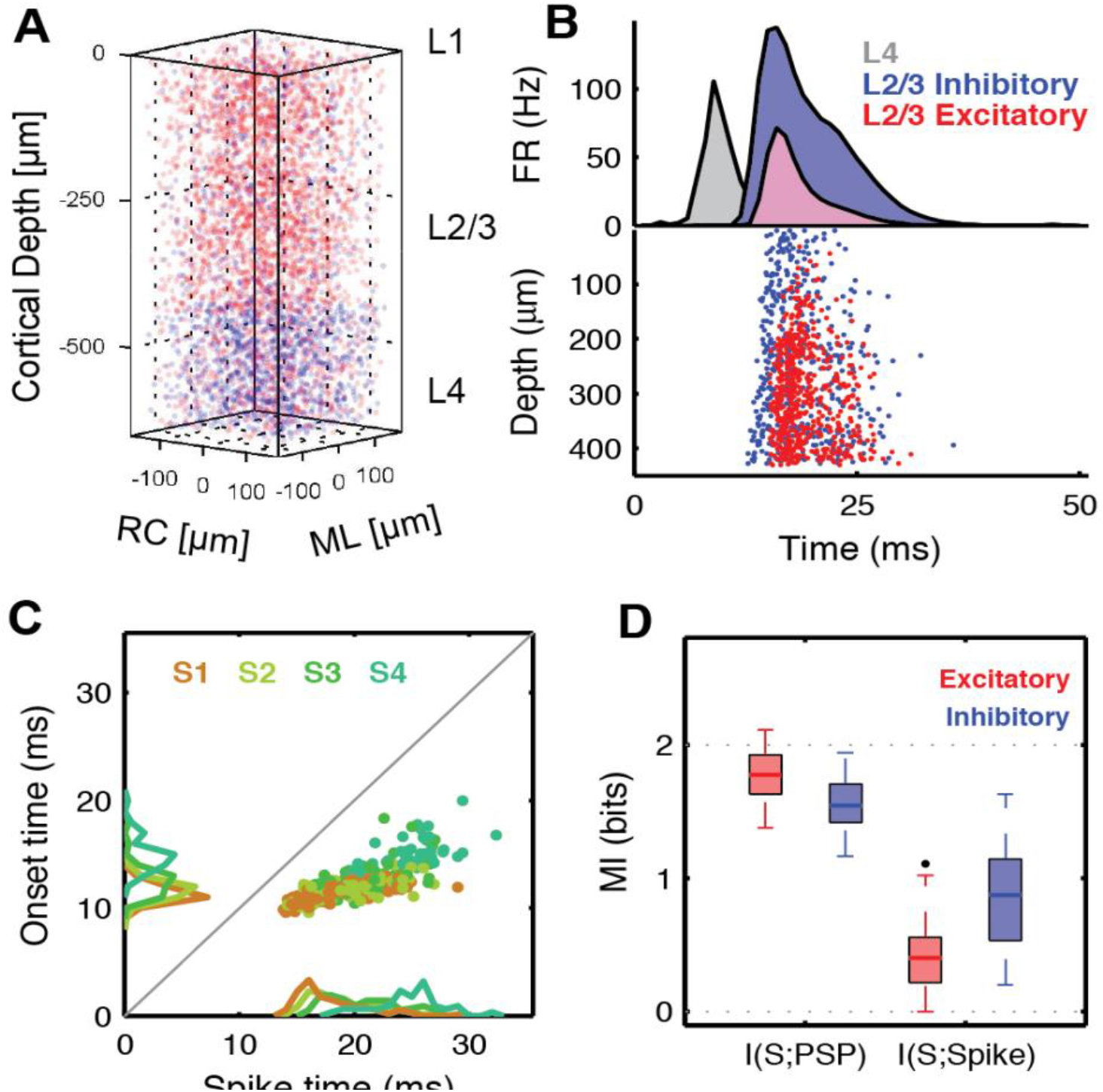
Anatomically constrained barrel column *in silico* reproduces the relationships between sub- and supra-threshold information. **(A)** An anatomically based model of a barrel column for L2/3/4 was generated to analyze the information transfer between L4 and L2/3 in analogy to the physiological recordings (see Huang et al. (2020) for details). **(B)** In response to stimulation in L4 with a whisker-like PSTH (grey), excitatory (red) and inhibitory (blue) cells respond in L2/3, with inhibitory activity eventually extinguishing the total activity in the network. **(C)** Corresponding to the *in vitro*/*in vivo* data, the timing of PSPs for a given stimulus is more precise than the spikes they evoke (compare to Fig. 1L). **(D)** The relationship between PSPs and spikes in terms of timing and reliability leads to single cell mutual information very similar to the recorded data (excitatory cells, compare to Fig.2A). Inhibitory cells (not recorded), show less information in their PSP response, but more information in the spikes (all properties combined for both cell-types). Dotted line: stimulus entropy.

**Figure 4.**
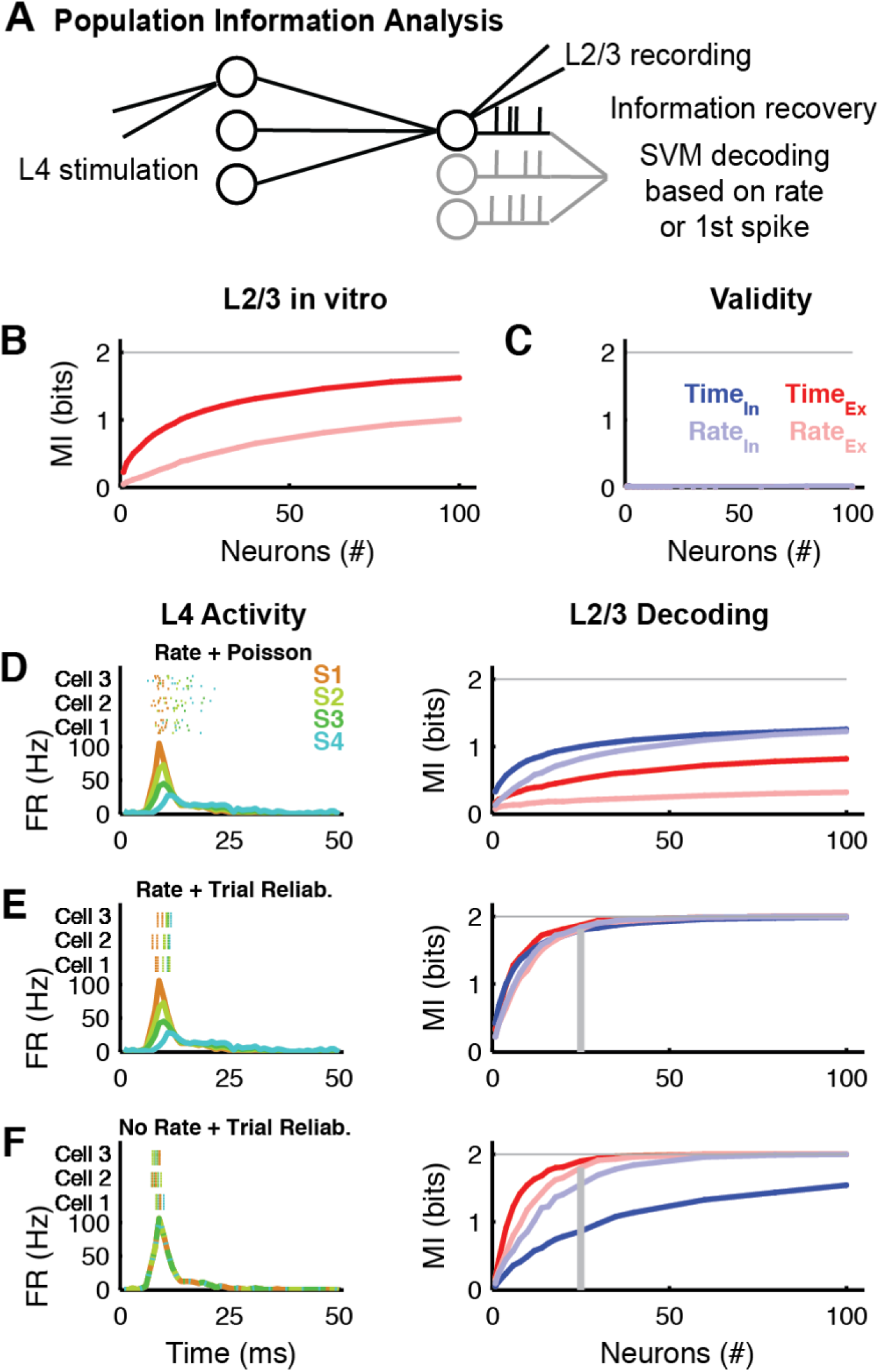
Information recovery in neural populations recorded *in vitro*. If a postsynaptic EPSP carries near complete information about the stimulus in the periphery (Figure 2), how does the postsynaptic neuron reconstruct this information from poorly informative action potentials of the presynaptic neurons? (A) To address this question we evaluate the mutual information from population spike trains of groups of excitatory or inhibitory neurons. To prevent the sampling bias, MI is estimated between the stimulus and an SVM decoding from the population response. (B) Population information estimated from bootstrapped *in vitro* recordings show nearly complete recovery of stimulus information. Asymptote is reached above 81% for 100 neurons for temporal decoding (dark red), and remains systematically lower for the rate-based decoding (light red). (C) Estimating population information for non-informative stimuli (identical PSTH, Poisson-spiking) leads to vanishingly low MI values, demonstrating that the analysis does not introduce a positive bias. (D) If the population activity in L4 is only constrained by the PSTH and otherwise spikes are drawn according to Poisson-distributions (bottom left, different colors = different stimuli), then inhibitory neurons carry more information for both time (dark blue) and rate (light blue) decoding, than excitatory neurons (dark & light red respectively). The gray line denotes the entropy of the stimulus. (E) If PSTHs differ across stimuli but spike timing is stereotypical across trials (‘Rate + Trial Reliability’, top left, multiple trials per neuron above each other), coding becomes highly effective and independent of the cell-type and coding strategy (~25 cells). (F) If L4 PSTHs do not distinguish stimuli, but only the timing of individual neurons across trials is stereotypical (No Rate + Trial Reliability, top left), a remarkable shift occurs, with excitatory neurons reaching almost complete information for much smaller group sizes (~25 cells). In all plots the vertical grey line indicates where 90% of the information is represented.

The model has anatomically correct numbers and laminar locations of major classes of inhibitory and excitatory neurons in L4 and L2/3 (Fig. 3A), single neuron dynamics based on experimental observations as well as statistically defined connectivity and synaptic transmission parameters. Stimulation was provided analogously to the *in vitro* stimulation in L4, using previously collected L4 peri-stimulus time histograms (PSTH) of principal and surround whisker stimulation *in vivo* (Fig. 3B, L4 response to principal whisker in gray (Celikel et al., 2004). PSTHs of simulated L2/3 excitatory and inhibitory neurons correspond to experimentally observed ones under similar conditions (De Kock et al., (2007), Fig. 3B red and blue, respectively).

In the model, the timing of and information in PSPs and spikes closely matched the properties of the real neurons in biological networks (compare Fig 3 to Fig.1-2). In the simulations, the trial-to-trial variability in spike timing was substantially and significantly larger than the variability in PSP timing (Fig. 3C, compare with Fig. 1A). Stimulus information was nearly fully retained in the somatic PSPs of excitatory neurons (Fig. 3D, red, 88.8% of the stimulus entropy), yet reduced substantially (20.1%) during spike generation, similar to our observations in biological neurons (Fig. 2A). Interestingly, inhibitory neurons carried significantly less information in their PSPs (Fig. 3D, blue, 77.3%), but also exhibited less information loss during spike generation (43.6%).

In the bootstrapped data, the stimulus information could be almost fully recovered from populations of excitatory L2/3 neurons recorded *in vitro* (>81.1%, Fig. 4B). The amount of information recovered was substantially greater for decoding including timing (81.1%, in timing of the 1st spike, binned at 2ms, 100 cells), than for rate based only decoding (50.5%, red vs. light red) and was largely independent of the population size (i.e. the MI saturates quickly as a function of population size). To avoid an overestimation of information from high-dimensional population data (due to the limited sampling bias), we first decoded the stimulus from single trial responses using a support vector machine (SVM) based decoder (Fig. 4A) before computing the MI (Ince et al., 2010b, 2010a; Quian Quiroga and Panzeri, 2009). To verify that this method did not introduce a positive bias, we computed the information in response to an artificial uninformative stimulus set (same PSTH for all stimuli, independent Poisson spiking), which yielded near-zero MI values (Fig. 4C). The performance of the SVM provided significantly better results (correctly predicting 94% of the stimuli), than linear (79%) or quadratic (80%) decoders. However, better decoders than the SVM may still exist, and the information calculated here therefore constitutes a lower bound on the available information in the population data. Also, it should be noted that the present timing code does not automatically include the rate code, since only the first spike is considered (see Materials and Methods).

#### Contribution of timing/rate reliability for information recovery

While a well-defined stimulus can be provided *in vitro*, the details of the L4 population activity cannot be controlled. From the perspective of information transmission in single trials, the most important property of the neural response is the reliability across trials. We utilized the barrel column *in silico* (Huang et al., 2020) to investigate the influence of the encoding strategy in L4 (reliability of spike timing and spike count) on the information transfer to L2/3.

We first considered three extreme cases of encoding in L4, with a more systematic exploration in the following section. In the first case (‘Rate + Poisson’), stimuli are encoded only by the population PSTH, but spikes across trials and neurons are drawn with Poisson statistics. In the second case (‘Rate + Trial Reliability’), stimuli are encoded by the population PSTH in L4, but also by the spike timing and count of its individual neurons (i.e. the spike trains were identical for each trial with the same stimulus conditions). Within the constraints of the experimentally observed PSTHs, these two cases constitute the lower and upper bound of trial-to-trial reliability in L4. In the third case (‘No Rate + Trial Reliability’), the population PSTH carries no information about the stimulus, but all information about the stimulus is encoded in the spike timing and count of individual neurons. The latter case is added for comparison with the other two encoding paradigms. Here, the population PSTH does not distinguish between stimuli, which happens for instance for texture recognition tasks (Arabzadeh et al., 2005). These three cases are illustrated in the insets of Fig. 4D-F. More details regarding the construction of these cases are given in Materials and Methods.

In the ‘Rate + Poisson’ case, information transfer is overall low, with interneurons providing a superior readout of the information about the stimulus in L2/3 compared to excitatory neurons, both when the information was decoded in rate and in timing (Fig 4D right, light and solid colors respectively, excitatory (red) vs. inhibitory (blue) neurons). While timing and rate codes are similarly efficient in interneurons, substantially larger populations of excitatory neurons are required to decode information in rate than in time. Given that the timing of the stimulus is only present on the population level in L4, the dominance of this temporal readout in L2/3 is remarkable. Assuming the emulated stimuli *in silico* approximate the stimuli *in vitro* with high accuracy (see Figs. 1 and 2), the ‘Rate + Poisson’ coding does not reflect the L4 encoding scheme in the present experiments.

In the ‘Rate + Trial Reliability’ case, the information transfer is overall substantially higher than in the ‘Rate+Poisson’ case (Fig. 4E right). In this condition the number of neurons required to recover the full stimulus information is the lowest (~25) of the three cases. This was expected, since in this case two sources of information - rate and timing - are used in the encoding of the stimulus. Remarkably, both cell-types and both decoding strategies yield very similar information values here, suggesting this encoding strategy is optimal for information transfer.

In the ‘No Rate + Trial Reliability’ case, the information transfer is intermediate between the two preceding cases. Here, the stimulus is only encoded by the responses of individual L4 neurons, not by the population PSTH. Interestingly, the opposite from the ‘Rate + Poisson’-case can be observed here in the decoding efficiency between L2/3 cell-types: contrary to the ‘Rate + Poisson’-case, where interneurons transfer more stimulus information, here excitatory neurons become substantially more efficient at representing information (compare Fig. 4D to 4F).

In summary, the availability of stimulus information in L2/3 spike trains is highly dependent on 1) the encoding properties of L4, 2) the decoding strategy in L2/3 and 3) the identity of the L2/3 neuronal populations (inhibitory or excitatory, for a summary see Table 1). Information about the stimulus in the spikes of L4 single units is best recovered by L2/3 excitatory neurons (Fig. 4F), given that there is a reasonable trial-to-trial reliability. Next, we systematically modulated the information content in L4 spike trains to investigate the consequences for L2/3 information availability.

**Table 1:**
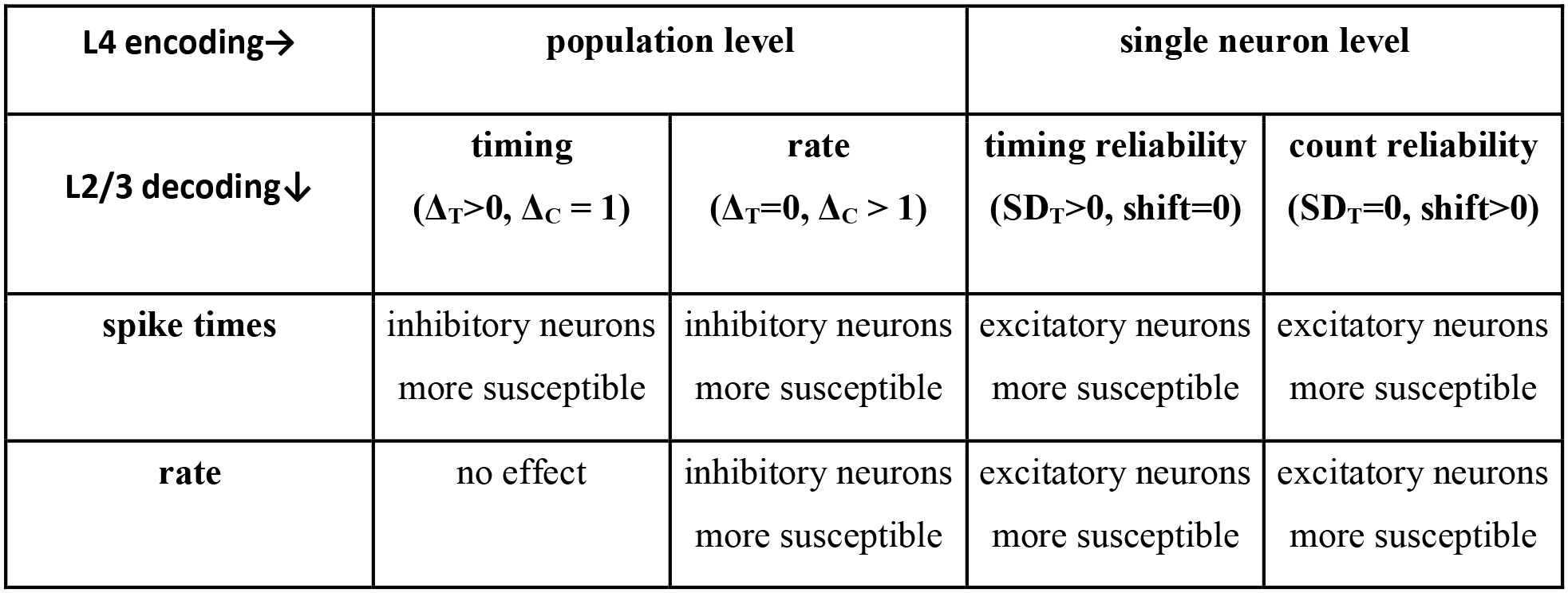
Summary of information recovery results depending on L4 encoding and L2/3 decoding schemes.

#### Population and single unit information selectively influence inhibitory or excitatory cells

While we only considered the extreme cases of L4 encoding above, realistic encoding will necessarily cover a range of cases between these extremes. Different stimuli will often, but not always, lead to different population PSTHs (but different surface textures may well lead to similar population PSTHs while finely modulating single unit responses (Arabzadeh et al., 2005). Conversely, even in cases where the population PSTH carries substantial information about the stimulus identity, spiking may well not be Poisson, but more reliable (especially at the response onset see e.g. (Amarasingham et al., 2006)).

We investigated the contributions from the population and the single unit separately. Information on the population level was represented as the average PSTH of the population (see Figure 6A1). Stimulus information was encoded in L4 spike trains either as timing or rate differences. Timing differences were implemented as shifts of the PSTHs (Δ_T_), whereas firing rate differences were implemented as rate factors between the PSTHs (Δ_C_). If Δ_T_ = 0 ms and Δ_C_ = 1, then no information is contained in the population PSTH. Conversely, if Δ_T_ = 4ms and Δ_C_ = 4, the combined differences between the PSTHs are similar to the experimentally observed ones. These parameters allow us to study the susceptibility of L2/3 neurons to the different encoding strategies of L4 neurons (examples of spike patterns are shown in Fig. 5A1 above the PSTHs, 10 trials each).

**Figure 5.**
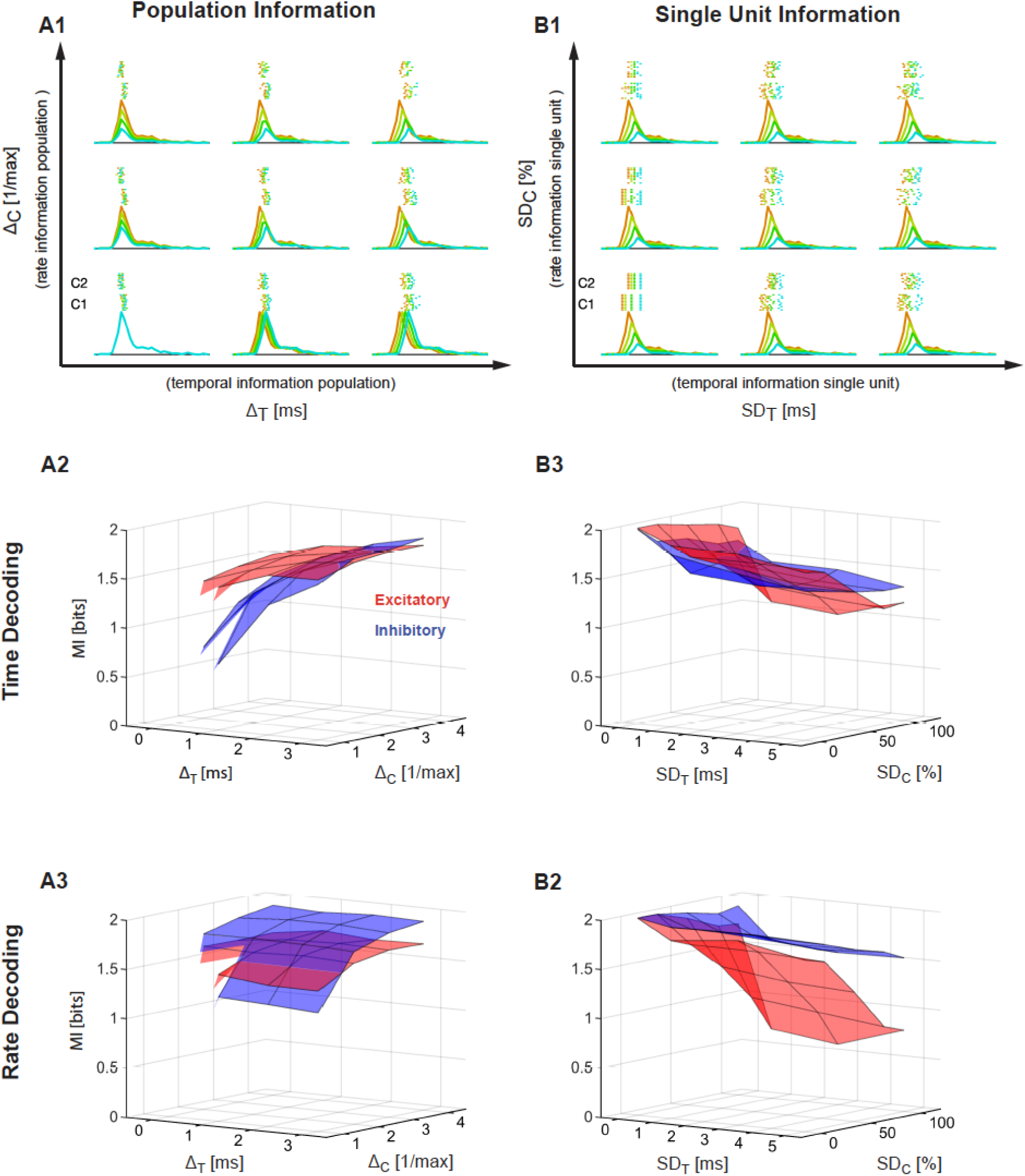
Stimulus encoding by presynaptic single neurons and populations of neurons selectively influences the decoding performance of the postsynaptic excitatory or inhibitory neurons, respectively. (A1) Stimulus information can be encoded in differences in rate or timing on the level of the population PSTH. Different combinations of these two coding dimensions are varied, with Δ_T_ (abscissa) indicating different timing for different stimuli (different colors, see Fig. 4), and Δ_C_ (ordinate) indicating different rates for different stimuli. Maximal information is achieved for high values of Δ_T_ and Δ_C_. For each condition the population PSTHs and two example cells are shown (raster plot for 10 trials, above). Spike-times of individual neurons are Poisson-distributed given the PSTH. NB C1 and C2 denote the responses of two different example cells. (A2) Decoding of first spike timing reveals a greater sensitivity of inhibitory neurons (blue) to the level of information in the L4 population response, both for time and rate information in L4. Conversely, excitatory neurons (red) are comparatively insensitive. (A3) Decoding of rate again reveals a greater sensitivity of inhibitory neurons to the level of information in the L4 response for different rates. Since we did not limit the time window of analysis, neither of the cell types is influenced by variation in time, while leaving the rate information unchanged. (B1) Stimulus information can also be encoded in the reliable discharge of single units. We modulated the reliability by introducing variability in timing (SD_T_) or variability in count (SD_C_) independent of each other. Maximal information is achieved for SD_T_ and SC_D_ both close to 0, i.e. perfectly reliable responses. Colors and raster plots as in A1. (B2) Decoding of first spike timing reveals a great sensitivity of excitatory neurons to the L4 information in single unit responses for both variability in time (SD_T_) and rate (SD_C_). (B3) Decoding of rate shows a very strong sensitivity of excitatory neurons on the single unit information. Conversely, inhibitory neurons exhibit almost no sensitivity to single unit information in L4, and are thus dominated by L4 population information.

**Figure 6.**
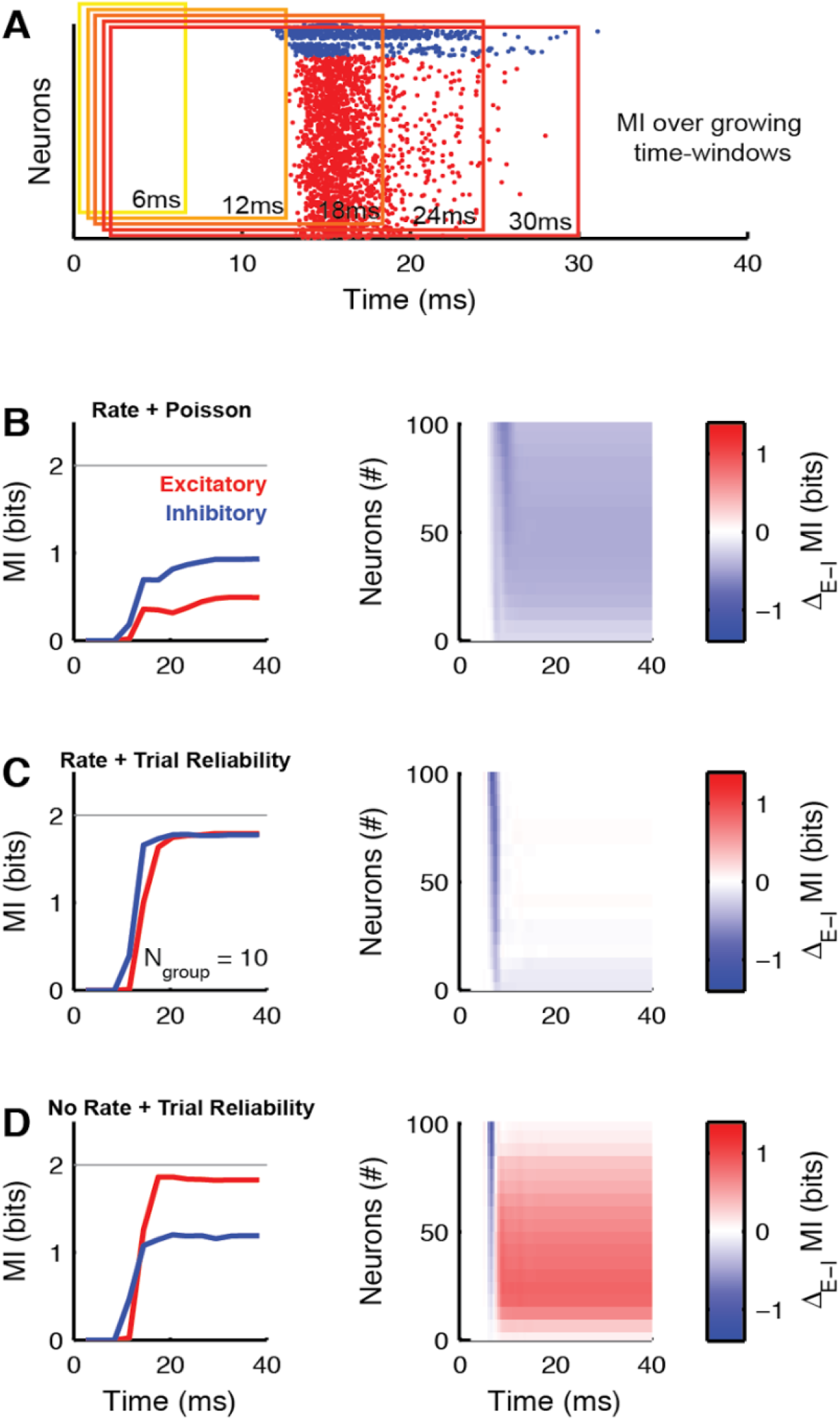
Neuronal information recovery is completed in <20ms after stimulus onset. **(A)** Time-scales of information recovery computed by calculating the MI between stimulus (not shown) and spike trains (single trial example shown) over time-windows of increasing lengths (6-30ms at 3ms steps) **(B)** For the ‘Rate + Poisson’ encoding in L4, both excitatory (red) and inhibitory (blue) neurons in L2/3 reach their respective maximal information (left: group size of 10 cells, the gray line denotes the entropy of the stimulus), ~25-30ms after stimulus onset. L2/3 inhibitory neurons (blue) encode more information, independent from the peri-stimulus time and group size (right). The color code shows the difference in MI between the excitatory and inhibitory groups, with red for larger MI in the excitatory, and blue for greater MI for the inhibitory neurons. **(C)** For the ‘Rate + Trial Reliability’ condition in L4, the information content of the two populations is quite similar (left) with a slight advantage for the inhibitory neurons at early times, but no dependence on group size (right). **(D)** In the ‘No Rate - Trial Reliability’’ case, the divergence between information content only begins around 12ms after stimulus onset, after which excitatory neurons achieve a substantial coding advantage, especially for smaller group sizes.

For decoding using spike times, inhibitory neurons exhibited a substantially greater susceptibility to variations in the distinguishability of stimuli on the L4 population level compared to excitatory neurons (blue vs. red, Fig. 5A2). This was true for both variations in time (Δ_T_) and rate (Δ_C_) in L4. Similarly, for rate decoding, inhibitory neurons were more susceptible to changes in rate than excitatory neurons (Fig. 5A3). Timing had little effect on rate decoding, since this corresponds mostly to a shift in the analysis window (unrestricted here), with no change in rate information in L2/3.

Next, we considered the influence of various levels of single unit information on the information availability in L2/3. Here, the population PSTH is kept fixed, but the temporal and count reliability are varied on a neuron-to-neuron basis (see Fig. 5B1). The timing reliability was varied by introducing a temporal jitter to individual spikes across trials (SD_T_), while contracting spiking patterns to remain consistent with the population PSTH. Count reliability was varied selectively by a linear transition between a completely reliable and a Poisson model, while maintaining the population PSTHs. This was done by shifting spikes between neurons while maintaining the temporal variability across neurons.

For decoding using spike times, excitatory neurons showed much greater susceptibility to single unit differences in reliability in L4, both for rate and time (Fig. 5B2). Interestingly, this carried over to the rate decoding to an even greater degree, which may be the domain of action for inhibitory neurons (Fig. 5B3).

In summary, L2/3 excitatory neurons are much more sensitive than L2/3 inhibitory neurons to the spike timing of L4 single neurons, whereas the information encoded by the population PSTH in L4 is carried mostly by inhibitory L2/3 neurons. Hence, we propose that the inhibitory and excitatory populations perform stimulus decoding in parallel, extracting stimulus information from distinct features in L4 activity (see Table 1). Together they have the ability to represent the entire information efficiently in small populations.

### Information recovery occurs rapidly within a few milliseconds

Information processing in the sensory cortices is under severe temporal constraints, especially in S1, where the sensory input is tightly integrated with the motor output for the purpose of precise and adaptive whisking control (Li et al., 2015; Proville et al., 2014; Voigts et al., 2015, 2008). The state of processing at a given time can be estimated by computing the mutual information over limited time windows, which progressively include a larger proportion of the neural response (Fig. 6A, all excitatory and inhibitory neurons separated for a single trial). In combination with varying the group size, we can thus obtain a ‘neurotemporal’ overview over the process of information availability in L2/3 as a function of neuronal class.

We consider three different encoding strategies by L4, ‘Rate + Poisson’, ‘Rate + Trial Reliability’ and ‘No Rate + Trial Reliability’ as in the previous section (Fig. 6B-D). In the ‘Rate + Poisson’ case, the mutual information begins to increase with interneurons leading over excitatory neurons (Fig. 6B, left, group size = 10 cells) around 12-14ms after stimulus onset (in L4). The inhibitory neurons reach maximal stimulus information, and do not achieve full stimulus information. Groups of inhibitory neurons encode more information than excitatory neurons, independent of the time relative to the stimulus onset and almost independent of group size (Fig. 6B, right).

For the ‘Rate + Trial Reliability’ encoding condition in L4, the difference in the information content between cell types in L2/3 is small, with inhibitory neurons carrying slightly more information at early peri-stimulus times and across all group-sizes (Fig. 6C, right). The difference in onset timing renders the information content of the inhibitory neurons higher only during the initial 1-2ms after response onset, due to the earlier response times of the inhibitory neurons (Fig. 6C, left).

In the ‘No Rate + Trial Reliability’ condition, the times when the information content increases are very comparable for excitatory and inhibitory neurons. However, after a few milliseconds, excitatory neurons prevail over inhibitory neurons. This advantage is preserved over time, whereas the difference in information content as a function of group size is strongly reduced, with inhibitory neurons eventually catching up with excitatory neurons (Fig. 6D, right).

In summary, the representation of stimulus information in L2/3 is rapidly completed within only a few milliseconds (3-5) after response onset. Which neurons, i.e. excitatory or inhibitory, carry more stimulus information is determined by the encoding strategy in L4 (corresponding potentially to different types of stimuli), but not much on the peri-stimulus time. As before, pure rate coding on the level of L4 is identified as an insufficient coding strategy, as it does not fit our experimental results of almost complete information recovery.

## Discussion

We demonstrated that although the intracellular information transfer, i.e. the PSP-to-action potential transformation, results in a significant loss of information about the stimulus, local networks can overcome this loss by integrating information from a small, experimentally tractable, number of neurons. Therefore, the somatic PSPs received by a single cortical neuron contain nearly complete information about the stimulus, even several synapses away from the sensory periphery. The efficiency of such information recovery is determined by a conjunction between the encoding scheme, neuronal class and decoding strategy. Excitatory and inhibitory cells take complementary roles in carrying information in single unit or population activity, respectively.

### Contribution of temporal coding in somatosensory cortex

Encoding information on short temporal scales can enrich the information content of neural activity relative to coarser average rates (Bialek et al., 1991; Bialek and Rieke, 1992). There has been a long discussion about whether the brain uses such a ‘spike code’ or ‘rate code’ (for a review, see Brette, 2015). It has been argued that since cortical networks are both noisy and very sensitive to perturbations, a rate code is the only way to perform reliable computations (London et al., 2010, but see Denève and Machens, 2016). However, others have pointed to the presence of temporally encoded information in the somatosensory (Alenda et al., 2010; Panzeri and Diamond, 2010; Petersen et al., 2001) and other sensory cortices (Kayser et al., 2012, 2010). In particular, the timing of the first (few) spike(s) in response to a stimulus conveys much of the information present in a spike train (Gollisch and Meister, 2008; Johansson and Birznieks, 2004). Consistently, we find that the majority of the information in the PSP is encoded in its timing. However, the timing of a spike in response to such a PSP is substantially more variable than the PSP timing, such that only a small proportion of the information in the PSP is transferred to the spike (Fig. 2). The amount of information loss could even be more substantial *in vivo* in the presence of background ongoing activity. On the population level, we find again that the temporal information is highly relevant during information recovery. In agreement with the previously observed importance of the first spikes, we find that the information content in the population asymptotes within 5 ms after the first spike in local populations, consistent with the time-scales of neuronal read-out in whisker cortex estimated before (5-8 ms, (Stüttgen and Schwarz, 2010)).

The temporal information described above can be fully characterized by single neuron variations in rate, and hence does not include higher order temporal codes, such as the pattern of inter-spike intervals. Due to the sparse response nature of supragranular excitatory neurons, such a fine-grained higher order temporal code could only exist in the inter-spike intervals of inhibitory neurons or spike-patterns across multiple (excitatory or inhibitory) neurons. The term temporal code is however still appropriate for our results, since the time scales of the response are not only reflecting dynamics in the stimulus, but correspond to intrinsic computations of the neural network (Nemenman et al., 2004).

For the present dataset, Shannon’s mutual information was computed with responses aligned to the stimulus onset. Recent work by Panzeri and colleagues (Panzeri et al., 2010; Panzeri and Diamond, 2010) have pointed out that such a reference time is not necessarily available to a decoder in S1. How would a change to an internal reference time, such as the efference copy of the whisking signal (Crochet et al., 2011; Crochet and Petersen, 2006; Poulet and Petersen, 2008) or a population-based timing (e.g. the “Columnar Synchronous Response”, CSR, event defined by (Panzeri and Diamond, 2010)) affect the present results? Assuming that the population response can be approximated by a set of individually recorded neurons (as in (Panzeri et al., 2010; Panzeri and Diamond, 2010)), the influence of such an intrinsic reference on our results would be only minor, since the relative timing - and thus the relative trial-to-trial variability in timing - would be the same as in the stimulus locked case. Hence, the information content would not be modified. If, on the other hand, synchronization between neural groups occurs, results could be significantly influenced, since then variability could be transferred from spikes to PSPs (in which case the alignment would be based on the near-synchronous CSR). According to Petersen and colleagues (Petersen et al., 2001), covariability, measured as noise correlation, was assessed to be ~0.1, and subsequent studies have found even lower values (Renart et al., 2010), suggesting that stimulus-independent synchronization is not substantial (note however the results of (Franke et al., 2016)

### High information availability and multiplexed codes

To understand ‘how the brain works’, we need to understand what the neural computations are that make an animal interact with its environment, i.e. how neural activity is transformed from the low-level response, to perceptual input, to the high-level neural activity that generates behavior (Eliasmith and Anderson, 2002). For instance, perceptual invariance (an object can be recognized as one and the same under different circumstances) and selectivity (an object can be distinguished from other, similar objects) need to be explained by any consistent theory of perception (Seung and Yuste, 2012). A model of how increasingly abstract features can be recognized by neural networks along the sensory axis was already explained by for instance the perceptron-model (Rosenblatt, 1958; Seung and Yuste, 2012). When neurons in each processing layer respond to only a single, increasingly abstract, preferred feature, they disregard necessarily a lot of information. Therefore, on the *single-neuron level*, the transformation from input to output is expected to be very sparse, and ‘lossy’. However, whether on a *population level* it is necessary to be able to fully reconstruct the stimulus, remains an open question. We have shown here that the entire stimulus information is maintained in layer 2/3 of the barrel cortex and encoded by local populations in a distributed fashion. This information can be recovered already on the basis of a small subset of neurons (~10-20, if single unit information is present) on short time-scales (~5ms relative to response onset), ensuring a lossless representation of the sensory world in real-time, i.e. before the next sensory information arrives from the periphery (in the case of active tactile exploration in freely behaving rodents, the inter-contact intervals are >30 ms (Voigts et al., 2015, 2008)). In different setups (Dalgleish et al., 2020), in mouse visual cortex (Sriram et al., 2020) and in salamander retina (Marre et al., 2015), comparable values have been reported. This suggests that the full stimulus information is needed for the computations at several levels. Combined with the single neuron selectivity, our results suggest that this network performs a form of coordinate transformation (Denève and Pouget, 2003). However, what the exact nature of the computations of this and downstream networks is, remains an open question.

The neural activity of excitatory neurons in cortical layers 2/3 is generally considered to be sparser than in Layer 4 (see (Barth and Poulet, 2012) for a review, although the evidence is not yet fully conclusive). This sparsity has been linked to higher selectivity of encoding, in terms of fewer, more specific features represented per neuron. This increased selectivity could be the reason for the observed information loss during the transformation of PSPs to spikes. It has been argued that the sparsity of the transformation of presynaptic spike trains to PSPs to postsynaptic spikes is the result of optimal non-linear processing: only redundant information, that has been gained before and can be predicted from previous activity, is discarded, and postsynaptic neurons only respond to ‘new’ information (Denève, 2008; Ujfalussy et al., 2015). Our result that the postsynaptic membrane potential still contains the full stimulus information, is in agreement with this argument, and the observation that most information is contained in the first spikes mentioned before could also be explained this way. However, whether the information that is lost in the spike-generating process is truly ‘discarded’ information, or, contrarily, redundant information, depends on the presumed decoding of the neuron: which information is redundant or essential depends on the message that needs to be conveyed.

The distributed persistence of complete stimulus information could provide a practical solution to one of the classical dilemmas of neural encoding: the compatibility between a specific feature and the context of the entire stimulus space. Concretely, a readout neuron in L2/3 may have privileged access to L4 neurons selective for one type of feature, with in addition access to a wide range of inputs from a random subset of the population. It could thus act as a comparator and evaluate the dominant feature in relation to a representation of the entire stimulus. This becomes especially relevant in the case of multiple concurrent stimulations on different whiskers, corresponding to the natural situation an animal is exposed to during active exploration (Voigts et al., 2015). In this case, multiple signals (i.e. the signals from multiple whiskers, that carry different spatial and temporal information) have to be processed by a single population of neurons. If this population can be separated into independent-subpopulations, this implies that a population consists of multiple channels (in the information-theory sense), but if this is not the case, multiple signals are coded by a single population, so the code becomes multiplexed: a single channel (population) carries complementary information through different codes. The observation that multiple subsets of neurons carry complete stimulus information and the observation that spike timing and firing rate of the same population (channel) can contain independent information about the stimulus hints at such multiplexing ((Panzeri et al., 2001; Quian Quiroga and Panzeri, 2009), for a review, see (Panzeri et al., 2010)). Our finding that different postsynaptic populations can decode the timing-encoded and rate-encoded information shows that the information from both coding schemes (rate and timing by inhibitory and excitatory neurons) of such multiplexed encoded information can also be used by the brain for further processing in later stages. Information in inhibitory populations can then be forwarded by for instance disynaptic (dis)inhibition and the modulation of firing rates or spike probabilities of excitatory populations. Multiplexed codes have been discussed recently in the context of local field oscillations (Alenda et al., 2010), and the presence of selective and general information as described herein may provide an additional example of multiplexing (Fig. 7).

**Figure 7.**
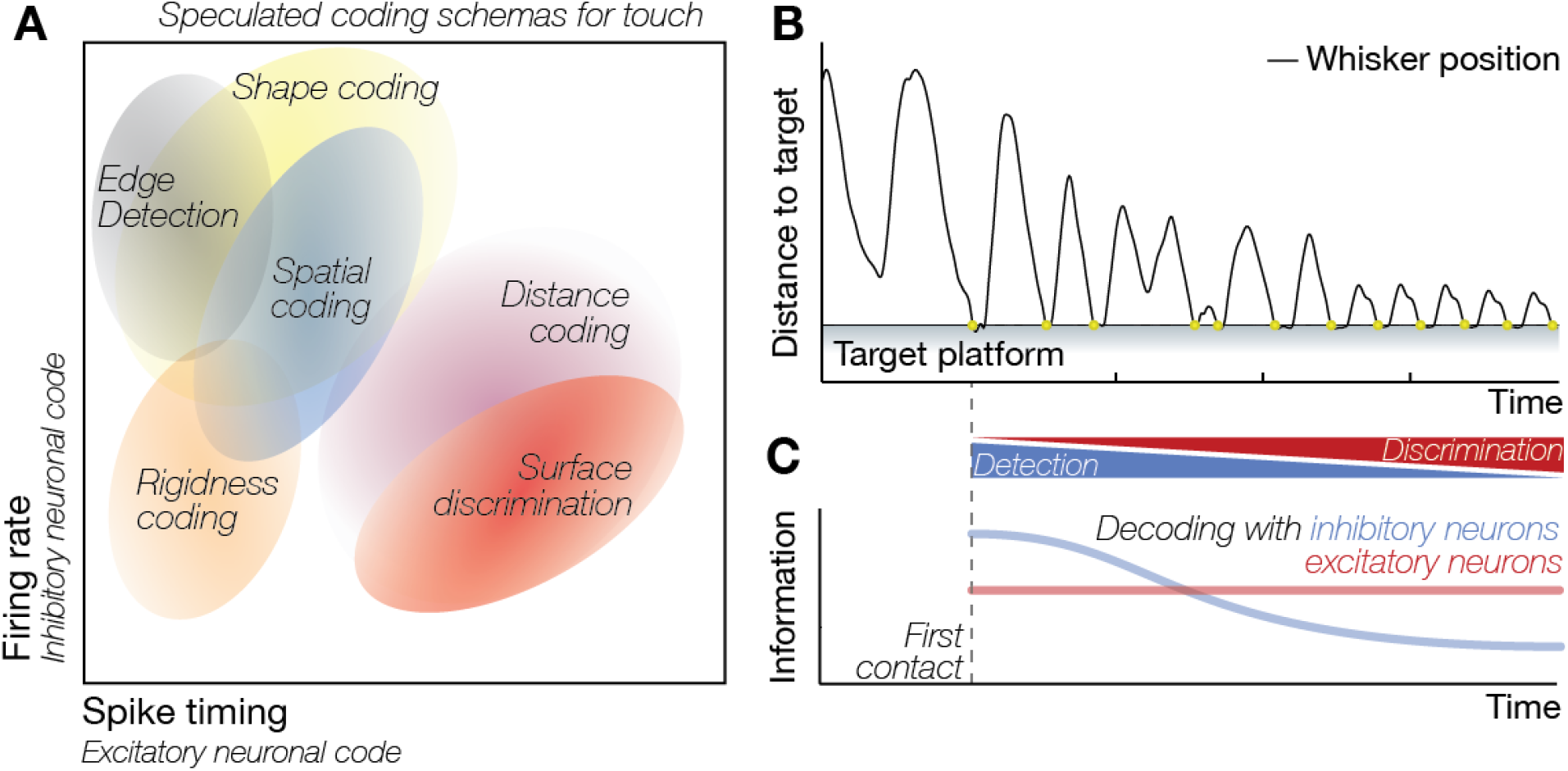
Multiplexed coding of touch. If intracellular information transfer, i.e. from EPSP-to-spike, results in a significant loss of near complete information, originally available in a single EPSP (Fig. 2), and if this information is recovered in local networks (Fig. 4 and 5) before the next sensory stimulus arrives (Fig. 6) using the rate and timing of spikes at the single cell and population levels (Fig. 4-6), selective decoding of stimulus properties by excitatory and inhibitory neural populations (Fig.5) will result in a multiplexed code for sensory processing. **(A)** If excitatory and inhibitory neurons preferentially decode the stimulus information from the spike timing of individual neurons and the population rate of presynaptic neuronal activity (Fig. 5), respectively, information content of the activity in excitatory and inhibitory neurons should vary predictably – see the suggested coding schema for touch. **(B-C)** The information content across the neural populations will also vary depending on the complexity of the stimulus. **(B)** During tactile object localization in freely behaving animals (Celikel and Sakmann, 2007; Voigts et al., 2015, 2008), for example, as the animal approaches the tactile target and makes multiple contacts, the information content will change not only because the kinematics of touch varies, e.g. the amplitude whisker deflections is reduced to match the predicted position of the sensory target (Voigts et al., 2015), but also the neurons will represent different features of the sensory target. **(C)** We speculate that information in the inhibitory neurons will better predict the stimulus location, although the information content of the excitatory neurons will eventually supersede as surface features are encoded with the subsequent contacts with the target.

Given the information content across the excitatory and inhibitory neural populations calculated herein, we speculate that distinct tactile features are encoded by rate and timing of spiking during information encoding and decoded by excitatory and inhibitory neurons separately (Fig. 7A). If an animal were to use its whiskers to locate a tactile target in space for example (Celikel and Sakmann, 2007; Peron et al., 2015), this model predicts that inhibitory neurons would carry the largest amount of information during the first contact as the animal detects the edge of the tactile target. Similarly, at the detection of a contact during passive whisking (Clem et al., 2008), inhibitory neurons would preferentially respond to the onset of touch, serving as an edge detector. The information content of different signals within the L4-to-L2/3 channel is temporally constrained as the animal continues to explore its immediate environment, and makes additional whisker contacts with the tactile target (Fig. 7B; (Voigts et al., 2015, 2008)), presumably to predict object distance and extract additional surface feature information about the target. With the change of whisking pattern, the sensory history and the statistics of the local network activity, relative information in the excitatory population will eventually dominate the neural representation of touch (Fig. 7C).

How to convey stimulus information both lossless and efficiently, and how this depends on physical properties such as network connectivity and node (neuron) properties, is an important open question in network science, and neuroscience specifically (Maheswaranathan et al., 2018; Mastrogiuseppe and Ostojic, 2017), in which computational models like the one we present here (Huang et al., 2020) play an invaluable role. Recently, it has been shown that there is a trade-off between the sparsity and the amount of recovered information in neural coding (Billings et al., 2014): lossless coding is only possible if the connectivity in the network is not too sparse. Specifically, the authors showed that optimal connectivity included only a few excitatory synapses and strong inhibition. Moreover, activity-dependent thresholds appear to play an invaluable role in such efficient information transmission (Billings et al., 2014; Huang et al., 2016).

For future *in vivo* studies, an important question will be, whether complete information representation persists if larger stimulus sets/spaces are considered, since it is expected that the dimensionality of the response, and hence the number of neurons needed for complete information recovery, depends on the complexity of the stimulus (Gao and Ganguli, 2015). Due to the requirements of accurate estimation of mutual information, we had to restrict the stimulus space to four stimuli in the context of whole-cell recordings (leading to ~300 trials per recorded cell). Note, however, that even under these conditions, trial-to-trial variability could have prevailed and prevented complete stimulus reconstruction on the single neuron and population level. Moreover, the present results can only provide a lower bound on the available information, since not all possible codes were explored and the decoding step between stimulus and response renders all results lower bounds (Quian Quiroga and Panzeri, 2009). In contrast to a previous study in the auditory cortex (Ince et al., 2013), we find that more complex decoding methods provide an improved decoding quality and hence more mutual information. Concretely, support vector machine decoding with radial basis functions provided superior performance (94%) than either diagonal linear (77%), linear (79%), or quadratic (80%) decoders. In order for the neural system to achieve this quality of decoding, it would, however, need to have readout mechanisms which use decoding strategies beyond linear or quadratic combinations.

### Predictions for cell-type specific coding strategies

The cortical population of neurons is composed of various cell-types, which differ in their morphology, location and physiology (De Kock et al., 2007; Narayanan et al., 2015; Oberlaender et al., 2012; Staiger et al., 2015). These differences suggest distinct roles in information processing, some of which have recently been elegantly elucidated (Ko et al., 2011). Coarsely, on the level of their firing patterns, inhibitory neurons can be distinguished from excitatory neurons, by more dense responses, based on a greater convergence of connections (reviewed in Harris and Mrsic-Flogel (2013)). The connectivity in the present model was set in precise accordance with the latest results from the literature from identified, pairwise recordings (see Huang et al. (2020) for detailed references), and consequently recreates these differences in firing behavior. Going beyond previous work, we find the coding balance to lean to either cell class, depending on the encoding strategy used in L4.

We explored these encoding strategies in L4, finding that excitatory neurons more effectively convey information encoded in L4 single units, requiring a level of reliability in L4 beyond Poisson-spiking (Figure 6B). Conversely, inhibitory neurons are more effective in carrying L4 population rate information (Figure 6A). Hence, together, excitatory and inhibitory neurons make effective use of the combined information in population rate and single unit responses in L4.

The L4 encoding is likely to depend on the stimulus condition: Many stimuli will induce time-varying population rates, which distinguish them from other stimuli. However, exceptions exist, such as the comparison of similar textures (Arabzadeh et al., 2005), which have only small differences in population rate, and differ more in their fine-structure. On the other hand, temporally structured inputs (e.g. many natural stimuli) lead to stronger time locking between neurons in L4 (Amarasingham et al., 2011; Litwin-Kumar and Doiron, 2012). Based on our results, we suggest that excitatory and inhibitory neurons might focus on distinct individual and population information to optimize the availability of stimulus information in local networks. Since long-range projections of inhibitory neurons are rare (Thomson and Lamy, 2007), the information content in the spiking of the inhibitory neurons is likely to be most relevant for local processing. Testing this hypothesis will not be trivial, since the inhibitory neurons cannot be removed from the network without influencing the overall network dynamics. Nonetheless, transient optogenetic modulation of the rate and timing of select inhibitory neurons’ activity while studying neural encoding of stimuli in the rest of the network will help to answer the question about which inhibitory neurons contribute more to the transfer and recovery of information within a column (i.e. local) and across columnar networks.

In summary, the results presented here suggest that single neurons are efficient real-time encoders of stimuli even several synapses away from the sensory periphery, but intracellular information transfer results in a substantial loss in the information transmitted to the postsynaptic neurons. The lost information can be recovered rapidly, i.e. within 20 ms, by comparatively small numbers of neurons in local populations, so that lossless information transfer along the sensory axis is ensured. The information recovery depends critically on the type of the neuron as well as the coding properties of both the presynaptic and postsynaptic pools of neurons, such that excitatory and inhibitory populations process complementary information about the stimulus in the somatosensory cortex.

## Materials and Methods

### Experimental procedures

Rats from either sex were used according to the Guidelines of National Institutes of Health and were approved by the local Institutional Animal Care and Use Committee. All data can be found in this online repository: https://doi.org/10.34973/59my-jm24, the relevant code can be found here: https://github.com/DepartmentofNeurophysiology/Information-transfer-and-recovery-for-sense-of-touch-code-for-figures

#### In vitro recordings

In vitro whole-cell current-clamp recordings were performed in acutely prepared slices of the barrel cortex between P18-21, after maturation of evoked neurotransmitter release (Martens et al., 2015) as described before (Allen et al., 2003; Celikel et al., 2004; Clem et al., 2008). Oblique thalamocortical slices (300 mm, (Finnerty and Connors, 2000)) were cut 45° from the midsagittal plane in chilled low-calcium, low-sodium Ringer’s solution (in mM; sucrose, 250; KCl, 2.5; MgSO_4_.7H_2_O, 4; NaH_2_PO_4_.H_2_O, 1; HEPES, 15; D-(+)-glucose, 11; CaCl_2_, 0.1). Slices were first incubated at 37°C for 45 minutes and were subsequently kept in room temperature in carbonated (95% O2/5% CO2) bath solution (pH 7.4, normal Ringer’s solution: in mM, NaCl, 119; KCl, 2.5; MgSO_4_, 1.3; NaH_2_PO_4_, 1; NaHCO_3_, 26.3; D-(+)-glucose, 11; CaCl_2_, 2.5).

Visualized whole-cell recordings were performed using an Axoclamp-2B amplifier under IR-DIC illumination. A custom-made tungsten bipolar extracellular stimulation electrode (inter-tip distance 150 micrometer) was placed in the lower half of a L4 barrel. Stimulation protocol was as described before (Huang et al., 2016). In short, 10 ms long current pulses were delivered using a bipolar electrode located in the lower half of a mystacial whisker’s barrel. The pulses were square and had equal maximal amplitude although the rising phase of the stimulus had different slopes. It took 0,2,4 or 6 ms for the pulse to reach the maximum amplitude for stimulus (S)1, S2, S3 and S4, respectively. All intracellular recordings (pipette resistance 3-4 MOhm) were performed in L2/3, orthogonal to the stimulation electrode within 150-300 μm of the cortical surface. The internal solution (pH 7.25) consisted of, in mM, potassium gluconate, 116; KCl, 6; NaCl, 2; HEPES, 20 mM; EGTA, 0.5; MgATP, 4; NaGTP, 0.3. For whole cell recordings, putative excitatory cells were selected based on pyramidal shaped somata, apical dendrites and distal tuft orientation, and regular pattern of spiking to somatic current injections (500 ms; data not shown). Data was low-pass filtered (2 kHz), digitized at 5 kHz using a 12-bit National Instruments data acquisition board and acquired using Strathclyde Electrophysiology Suite for offline data analysis.

#### In vivo recordings

In vivo whole-cell current-clamp recordings were performed under ketamine/xylazine anesthesia at P28-30. Anesthesia was induced using 100 mg/kg (ketamine) and 10 mg/kg (xylazine) mixture and maintained with intraperitoneal ketamine-only injections (20% of the initial dose) as necessary. Upon complete loss of facial and hind-limb motor reflexes, the skull was exposed. A head-bolt was fixed posterior to lambda using cyanoacrylate and was used to immobilize the animal during experiments.

The surface over the primary somatosensory cortex (from Bregma, −0.5mm to −2.5mm, from Midline −2.5mm to −4.5mm was thinned using a dental drill. The surface was kept moist with a thin layer of low-viscosity mineral oil to maintain the transparency of the thinned skull. Cortical representation of the D2 whisker was localized in the contralateral hemisphere using intrinsic optical imaging as described before (Stewart et al., 2013) while deflecting individual whiskers using piezoelectric actuators as described elsewhere (Celikel et al., 2004). The skull above the center of mass of the functional whisker representation was punctured using a 28 gauge needle to allow patch electrodes to access the cortical region of interest. All electrode penetrations were perpendicular to the cortical surface. In vivo whole-cell recordings were performed as described before (Margrie et al., 2002) with recording electrodes (6-7 MOhm) filled with the same intracellular solution used in slice experiments. Two different whisker deflection protocols were used: During optical mapping experiments single whiskers were deflected along the dorsoventral axis at 5 Hz with 8° deflections for 20 times with an inter-trial interval of 20 sec (Stewart et al., 2013). During electrophysiological recordings single dorsoventral whisker deflections were delivered at 0.2 Hz for 200 times. In each trial 4° whisker deflections were delivered at 10 Hz for 1s. Throughout the experiment the animal’s core body temperature was maintained at 36.5±0.5°C.

### Data analysis

All analyses were performed off-line in Matlab (Mathworks, Inc), the code for the figures can be found online: https://github.com/DepartmentofNeurophysiology/Information-transfer-and-recovery-for-sense-of-touch-code-for-figures. Raw voltage traces were smoothed using running window averaging (1ms window size) and the following variables were calculated for all evoked responses: Onset time (Ot, in ms): Latency of the postsynaptic potential (PSP) onset in respect to onset of the stimulus; Rise time (Rt, in ms): Time it takes for the membrane to reach 90% of the PSP amplitude relative to the onset of PSP; PSP slope (Sl, in mV/ms) between 10-90% of the PSP amplitude and amplitude of the EPSP (Amp, in mV). If the trial included an action potential, the peak of the EPSP was set to the spike threshold (Vt). The spike threshold was defined as the membrane potential value at which the second derivative of the membrane potential reached a maximum as described before (Wilent and Contreras, 2004). In slice recordings, resting membrane potential (Vm, in mV) was calculated as the average membrane potential in a 40 ms time window prior to the stimulus onset. For *in vivo* recordings the same time window was used but the sweep was included in the data analysis only if the variance of the membrane potential was < 0.5mV during the time window. For those sweeps in which a spike was observed, the spike threshold and spike latency (St) were also calculated.

#### Mutual information analysis for single neurons

Only cells with more than 250 acceptable sweeps (summed across all stimulus conditions) were used to perform Shannon information analysis. The mutual information (MI) between any two variables *S, R* can be calculated as

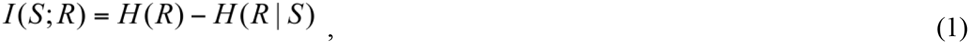

in which H is the entropy of a given variable R:

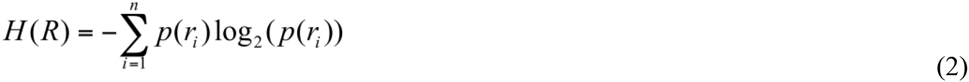

and H(R|S) is defined as

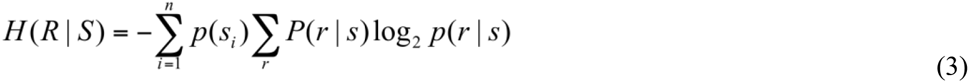

where i ranges over the stimulus/response types. Note that the stimulus entropy shows a small variability due to rejected trials. Similarly, the mutual information between one variable S and multiple **R** (joint mutual information) can also be calculated using equation (1). In this case, the synergistic effect of **R** can be expressed as the difference between the linear sum of the mutual information between S and each individual R and the joint information I(S;**R**):

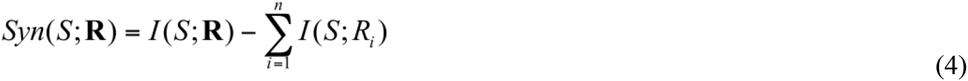

Information calculations were performed using the Information Breakdown toolbox (Magri et al., 2009) in Matlab (Mathworks. Inc). In short, each variable was first digitized using the equal space (‘eqspace’) binning method with 7 bins. The effect of different binning methods as well as the number of bins on MI values are also explored (Fig. S1). In the analysis based on the ‘eqpop’ binning method, the size of individual bins was modified so that a roughly equal number of observations was placed in each bin, instead of keeping the size of individual bins constant. Because in most trials only one spike was observed, only the first spike was considered when calculating the information in St. Thus, the spike latency St can be digitized to a single word, which has (number of bin + 1) possible outcomes, instead of a binary list which could have 2^(number of bin) possible values. Shuffle correction combined with Panzeri-Treves (Panzeri and Treves, 1996) bias correction was used to perform all information calculations for neural recordings (note that shuffle corrections can introduce a small source of variability, which can be seen for instance in comparing I(S,PSP) in Figure 2A with and H(S) or with the joint MI in 2C, or that can lead to an error bar above the stimulus entropy or below 0). The performance of the algorithm was evaluated by randomly selecting a subset of trials to calculate the mutual information (I(S;PSP), I(PSP;Vt) and I(PSP;St)), and subsequently checking the number of trials (Ns) needed for the calculated information values to reach asymptote (Fig. S2). When the ‘eqspace’ binning method was used, all information values reached asymptote after Ns > 70, well below the average Ns in the present data set (124±33.2 (range: 78-220) stimulus repetitions per stimulus).

#### Calculation of minimum observation size

An essential step in the information calculation method listed above is the estimation of the stimulus-response probability distributions from the experimental data. Following Panzeri and colleagues (Ince et al., 2010b) we calculated the number of experimental trials per stimulus condition, Ns, to be ~32 times larger than the number of possible response pattern, R, to get an accurate estimation (Ns/R≈32). This also means that to accurately estimate information between the subthreshold responses (Am, Sl, Ot, all binned to 7 bins) and the stimulus, 32×7×7×7 = 10976 trials/ stimulus =91h continuous recordings will be needed. Given technical infeasibility of maintaining whole cell access for the designated period we performed bias corrections to account for the upward bias in information estimation with limited sample sizes (see (Ince et al., 2010b) and (Victor, 2009) for further discussion). Methods like quadratic extrapolation (QE), Panzeri-Treves (PT) correction (Panzeri and Treves, 1996) and Nemenman-Shafee-Bialek (NSB) t experiments and 94±25.6 (range, 60-146) trials/stimulus for the *in vivo* whole-cell recordings.

#### Mutual information analysis for multiple neurons

For multi-neuron MI analysis we followed the approach to first decode and then estimate the MI between stimuli and the confusion matrix of the decode (Ince et al., 2010b; Panzeri and Diamond, 2010; Quian Quiroga and Panzeri, 2009) using support vector machine (SVM) in MATLAB with radial basis functions as the kernel transform. We utilized 90/10% cross validation during decoding to obtain an estimate of the generalized performance of the decoder. SVM decoding outperformed other decoders with an average performance of 94%, compared with some other decoders (diagonal linear (77%), linear (79%), quadratic (80%)). The use of an intermediate decoder ensured that the calculation was bias free (given that we observe the correct value of 0 bits for an uninformative set of stimuli, with otherwise very similar properties (see Fig. 3G)), but came at the expense of lower bound in MI estimates since a (potentially existing) better decoder would improve the MI.

For the in-vitro recordings we first had to generate bootstrapped populations of sufficient size to perform the population MI calculations. In order to preserve the within-cell variability of responses across stimulus and trials, we only drew bootstrap samples from the trials of each cell independently. As in the simulations we drew 100 samples of groups of each population size. Curves in Fig. 4 display averages over these samples.

### Network Simulations

The reconstruction of information in the neural network was performed in an *in silico* model of the barrel cortex.

#### Neural network

The model included a realistic account of the number of (Izhikevich, 2004, 2003) neurons and connectivity (Supplemental Table 1) within a barrel column for Layers 2/3, with inputs arriving from the L4, mimicking the conditions in the *in vitro*/*in vivo* experiments. For more details on the network model, see (Huang et al., 2020).

Synaptic currents in this network were modeled by a double-exponential function. Parameters of those functions (peak amplitude, rise time, half width, and pair-pulse ratio) were adjusted to match experimentally measured PSPs in barrel cortex (Supplemental Table 1; see Thomson and Lamy (2007) for a review). The onset latency was calculated from the distance between cell pairs; the conduction velocity of action potential was set to 190μm/ms.

Differences in activation state of cortex were included in the model by setting the common initial voltage and the equilibrium potential ***vr*** of all cells to −80, −70, or −60mV in a third of the trials, thus accounting for potential up- and down-states as well as an intermediate state.

#### Synaptic input from layer 4

Layer 4 stimulation was provided in the model based on population PSTHs collected extracellularly in anesthetized animals *in vivo* (Celikel et al., 2004). We used PSTHs of principal and 1st order surround whisker stimulation, as well as two linear interpolations between the two, yielding 4 stimuli with 2 bits total entropy, matching the numbers in the *in vitro* experiments. The PSTHs only specified the population firing rate in L4. We further explored population coding properties, by modifying the variability of spike timing across trials. If response times and spike counts were conserved across multiple trials, spike timing and counts within and between neurons start to carry additional information.

In the ‘Rate + Poisson’ condition, we assumed no trial-to-trial reliability beyond that given by the PSTH. Spike times were drawn based on Poisson statistics for each time with the PSTH modulating the firing rate (see Fig. 4D left). This condition forms a lower bound on the transferred information between L4 and L2/3, under the experimental constraints on the model. On the other extreme, in the ‘Rate + Trial Reliability’ condition, the PSTHs varied as before, but in addition neurons emitted the same sequence of spikes for every trial, preserving timing and count perfectly. This condition forms an upper bound on the information transfer, since within the experimental constraints no additional variability is introduced, which would reduce the mutual information. Finally, we consider the ‘No Rate + Trial Reliability’ condition, where the population PSTHs are uninformative across stimuli, and stimulus information is only contained in the spike trains of individual neurons. This case is a reference for other stimulus scenarios, where the PSTH may not vary much (e.g. texture-type stimuli), and individual timing becomes more important.

We also explored conditions between these extremes (Fig. 5), where the information in population or single neuron response was systematically varied. For the case of the population response we varied the different in time and firing rate of the PSTHs for different stimuli (Fig. 5A). Time differences were implemented by simply shifting the entire PSTH in time (tested shifts: [0,1,2,3] ms per stimulus, i.e. maximum shift was 9 ms). Rate differences were implemented as the fraction between the maximal and the minimal stimulus (tested fractions were [1,2,3,4], where e.g. 4 corresponds to the weakest stimulus being 25% of the strongest stimulus at the peak of the PSTH). The case of time shift 0 ms and rate fraction 1 is uninformative on the level of population rate. Single neuron reliability in this case was chosen as a medium level of single unit reliability (SD_T_ = 3 ms, SD_C_ = 20%). Single neuron reliability in response was also explored in timing and rate (Fig. 5B). Starting from perfect timing and rate, we degraded the information extractable from single neurons, by introducing timing variability (spike times were shifted by Gaussian-distributed noise with standard deviation SD_T_) and rate/count variability (spikes were deleted or added, by linearly mixing between Poisson and perfectly reliable spiking, with mixing parameter SD_C_, denoted as % in the figure). For both procedures, the modifications were performed while keeping the population PSTH approximately unchanged, i.e. for timing the overall timing distribution was contracted to keep the original PSTH, and for rate, spikes were shifted between neurons, rather than only removed from individual neurons.

These independent variations of population and single unit responses allowed us to separate the contribution of these two information sources to the information available in groups of L2/3 excitatory and inhibitory neurons (see Results).

## Abbreviations

AP: Action potential / spike
PW: Principal whisker
H: Entropy
S: Stimulus
Δ_T_: Variation in spike timing
L: (Cortical) layer
PSP: Postsynaptic potential
SW: Surround whisker
I: (Mutual) Information
PSTH: Peristimulus time histogram
Δ_C_: Variation in spike rate
CSR: Columnar Synchronous Response

## Acknowledgements

We are grateful to Drs. Rasmus Petersen and Alessandro Treves for their comments on a previous version of this manuscript, and the members of the Department of Neurophysiology for critical discussions. This work was supported by the Whitehall Foundation, Alfred P. Sloan Foundation, European Commission (Horizon2020, nr. 660328) and Netherlands Organisation for Scientific Research (NWO-ALW Open Competition; nr. 824.14.022 and NWO Veni Research Grant, nr. 863.150.25 to FZ) and a Christine Mohrmann Grant to FZ. The funders had no role in study design, data collection and analysis, decision to publish, or preparation of the manuscript.

