## Supplemental Figures and Tables for "Information transfer and recovery for the sense of touch"

### Supplemental Materials

##### Supplemental table

| Post vs Presynaptic | Input layer neuron | Excitatory neuron | FS interneuron | NFS interneuron |
|---|---|---|---|---|
| **Excitatory neuron** | | | | |
| **Pconn** | 0.11 | 0.03 | 0.14 | 0.10 |
| **uPSP strength, mV** | 0.7±0.6 | 1.0±0.7 | -2.96±2.52 | -0.49±0.49 |
| **uPSP rise time, ms** | 0.8±0.3 | 0.7±0.2 | 6.5±3.7 | 5.4±2.2 |
| **uPSP half width, ms** | 12.7±3.5 | 15.7±4.5 | 55.9±22.6 | 56.2±24.1 |
| **PPR** | 0.90±0.39 | 0.61±0.41 | 0.46±0.17 | 0.91±0.56 |
| **FS interneuron** | | | | |
| **Pconn** | 0.20 | 0.21 | 0.06 | 0.09 |
| **uPSP strength, mV** | 0.96±0.93 | 3.48±2.52 | -2.96±2.52 | -0.37±0.33 |
| **uPSP rise time, ms** | 0.89±0.31 | 2.32±1.00 | 6.5±3.7 | 3.1±2.0 |
| **uPSP half width, ms** | 15.0±8.2 | 16.3±5.8 | 55.9±22.6 | 20.0±12.1 |
| **PPR** | 0.99±0.66 | 0.70±0.14 | 0.46±0.17 | 0.91±0.56 |
| **NFS interneuron** | | | | |
| **Pconn** | 0.20 | 0.14 | 0.07 | 0.11 |
| **uPSP strength, mV** | 1.2±0.9 | 1.36±0.78 | -2.96±2.52 | -0.49±0.56 |
| **uPSP rise time, ms** | 0.42±0.1 | 1.3±0.5 | 6.5±3.7 | 4.9±5.4 |
| **uPSP half width, ms** | 14.0±9.1 | 14.0±6.2 | 55.9±22.6 | 33.3±12.0 |
| **PPR** | 1.00±0.62 | 0.96±0.51 | 0.46±0.17 | 0.91±0.56 |

**Supplemental Table 1. Connection probability and unitary PSP parameters between different neuron populations.**

Pconn, connection probability; uPSP, unitary postsynaptic potential; PPR, pair-pulse ratio. Values are mean±std. Connectivity was determined using axonal and dendritic projection patterns which were approximated by 2-D Gaussian functions. Synapses were distributed between axonal projections of the presynaptic neurons and dendrites of the postsynaptic neurons. This resulted in a binary connectivity matrix which determines the map of connections between any given neurons in the network. The connection probability between different types of neurons matched experimental observations (Avermann et al., 2012; Blatow et al., 2003; Caputi et al., 2009; Feldmeyer et al., 2006, 2002; Helmstaedter et al., 2008; Holmgren et al., 2003; Lübke et al., 2003).

**Supplemental Figures**

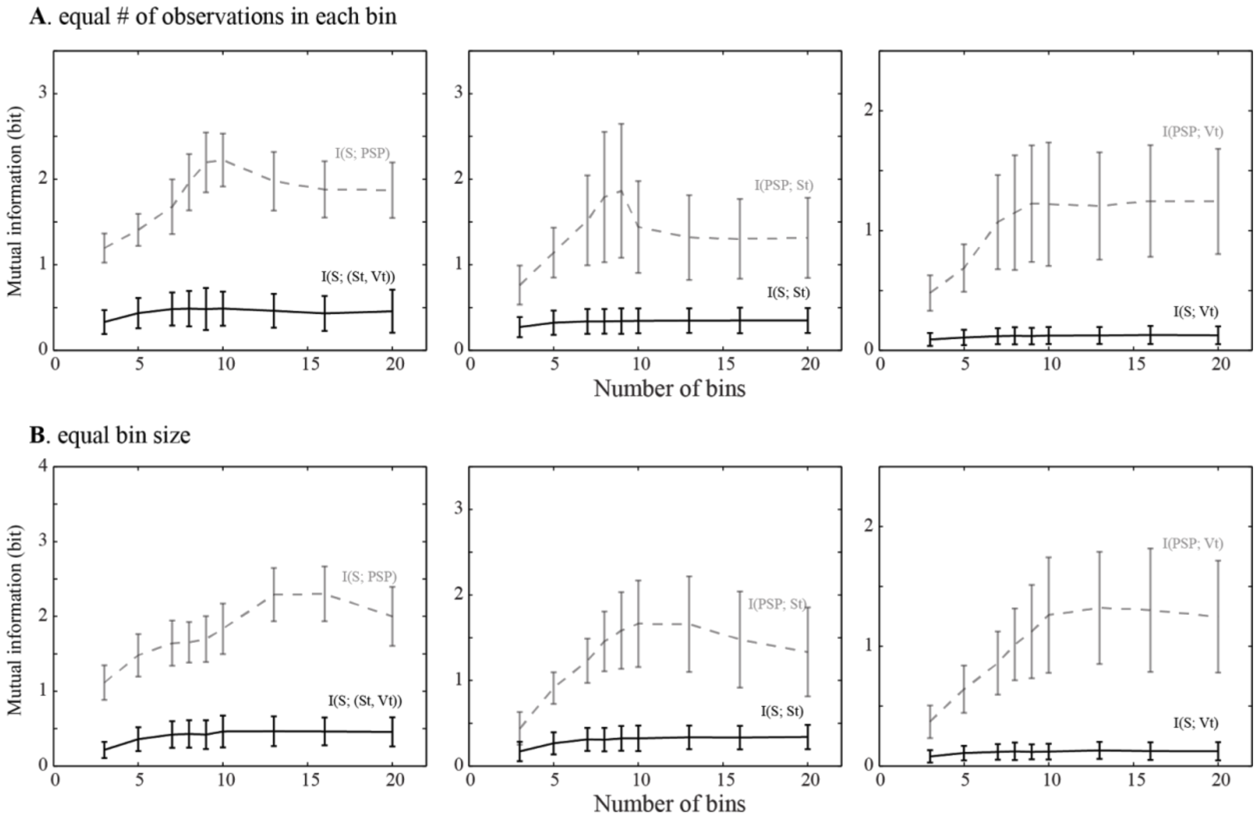

**Supplemental Figure 1. Effect of the bin number on mutual information calculations.** Two different binning methods were considered: **(A)** equal number of observations in each bin and **(B)** equal bin size. Independent of the bin number used in the calculations reported in the main text, PSP channels contain significantly more information about Stimulus (or the suprathreshold response) compared to action potential threshold (Vt) or spike timing (St).

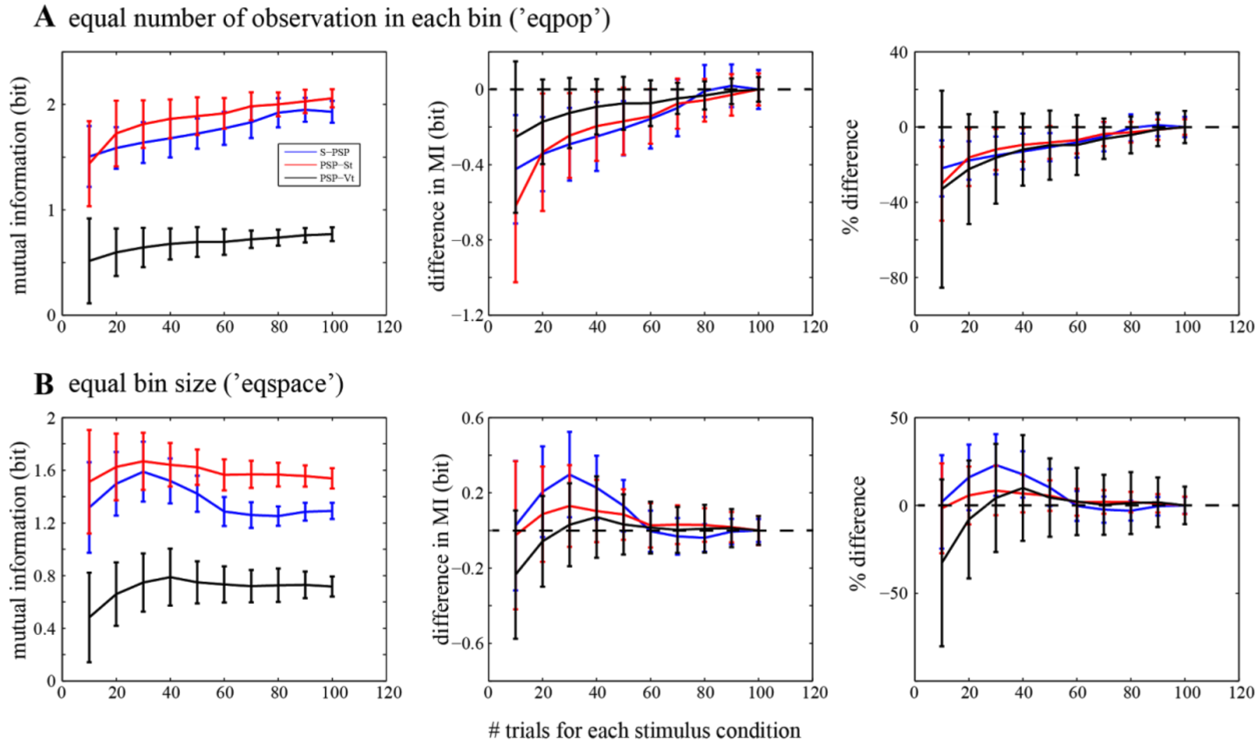

**Supplemental Figure 2. Change in mutual information as a function of the number of trials in each stimulus condition.** A random subset of trials in each stimulus condition was drawn to perform calculation on I(S; PSP), I(PSP; St) and I(PSP; Vt). **A)** Analysis based on 'eqpop' binning method (see Materials and Methods), in which the size of individual bins was modified so roughly equal number of observations was placed in each bin. **B)** Analysis based on 'eqspace' binning method (see Materials and Methods), in which the size of individual bins was kept constant. All three information values near the asymptote after ~70 trials in each stimulus condition. Therefore calculations in the main text were performed using >70 trials per condition (78-220, on average 124±33.2 trials/stimulus). Dashed lines denote a vanishing difference, compared with the information value calculated with 100 trials per stimulus condition.
